# Gene function predicts divergence across molecular levels

**DOI:** 10.64898/2026.01.13.699300

**Authors:** Antara Anika Piya, Ksenia Macias Calix, Raquel Assis

**Author notes:** Corresponding author:* Raquel Assis (R.A.).

## Abstract

Genes can evolve through changes at their DNA, RNA, or protein levels. However, because these changes are measured using distinct and often incomparable metrics, their relative contributions to genic evolution remain unclear. Here, we address this challenge by developing a standardized framework for comparing evolutionary divergence in protein-coding sequences, multi-tissue expression profiles, and protein structures. Application of this approach to data from *Drosophila*, rodents, primates, and *Arabidopsis* reveals substantial variation in divergence patterns across taxa, with only sequence divergence matching expectations based on evolutionary theory. Moreover, sequences evolve slowest and protein structures fastest in all taxa, consistent with the idea that higher levels of biological organization are closer proxies for the functions on which selection acts. Yet, weak correlations among divergence measures suggest that targets of selection vary across genes, perhaps depending on their functions. Indeed, few genes exhibit similar evolutionary patterns across all three levels, and different functions are enriched in genes with low or high divergence in protein-coding sequences, gene expression profiles, and protein structures. Together, these findings support the hypothesis that evolutionary targets of genes depend on their functions, shedding light on how selection shapes different levels of biological organization across taxa.

## Introduction

Evolutionary divergence is often used to identify genes under strong constraint due to their essential biological functions [Kimura, 1968, Consortium, 2004, Jordan et al., 2005, Brawand et al., 2011, Waterhouse et al., 2011, Warnefors and Kaessmann, 2013] or under weak constraint due to their roles in adaptation [Kimura, 1968, Barbash et al., 2004, Jordan et al., 2005, Cai and Petrov, 2010, Brawand et al., 2011, Assis, 2019a, Piya et al., 2023]. While early studies primarily focused on gene sequences [Kimura, 1968, Gummerson and Williamson, 1974, Miyata et al., 1980, Zurawski et al., 1984, Kimura, 1985, Wu and Li, 1985, Yang and Nielsen, 2000], the recent availability of diverse trait data has enabled genome-wide explorations of other classes of evolutionary divergence, including those corresponding to gene expression profiles [Jordan et al., 2005, Consortium, 2004, Brawand et al., 2011, Guo et al., 2016, Assis, 2019a, Piya et al., 2023] and protein structures [Lecomte et al., 2005, Williams and Lovell, 2009, Borcherds et al., 2013, Bordin et al., 2021]. Because natural selection acts on phenotypes, which are often difficult to link to their underlying sequence changes, analyses of expression or structural divergence may offer more direct insights into selective pressures and adaptive processes [Carroll, 2005, Gilad et al., 2006, Dean and Thornton, 2007, Fraser et al., 2011] Moreover, key questions remain about how evolutionary divergence varies across taxa and molecular levels, and what such differences reveal about genic targets of selection.

Findings from numerous studies indicate widespread variation in evolutionary divergence across both taxa and molecular levels. Specifically, taxon-specific differences in evolutionary rates have been observed in protein-coding sequences, gene expression profiles, and protein structures [Lemos et al., 2005, Jordan et al., 2005, Gillis and Pavlidis, 2009, Buschiazzo et al., 2012, Warnefors and Kaessmann, 2013, Konate et al., 2019, James et al., 2021, Lau et al., 2024]. Additionally, comparisons of coding sequence and expression divergence have uncovered weak-to-moderate correlations [Nuzhdin et al., 2004, Khaitovich et al., 2005, Lemos et al., 2005, Tirosh and Barkai, 2008, Kalinka et al., 2010, Assis and Kondrashov, 2014, Harnqvist et al., 2021, Zhong et al., 2021]. While this imperfect relationship supports the hypothesis that regulatory changes play a prominent role in expression evolution [Carroll, 2008, Emerson and Li, 2010], it also implies that coding sequence and expression divergence for the same gene often differ, potentially leading to conflicting evolutionary conclusions. Similarly, analyses of coding sequence and protein structural divergence have identified variable and complex associations[Wood and Pearson, 1999, Krissinel, 2007]. Although the relationship between gene expression and protein structural divergence remains elusive, with studies reporting both weak positive and weak negative associations [Paliy et al., 2008, Singh and Dash, 2008], the prevailing consensus is that their correlation is weak. Together, these findings suggest that different classes of evolutionary divergence capture distinct yet interconnected aspects of molecular evolution, highlighting the need for further investigation into the mechanisms driving their variation across taxa and molecular levels.

Despite these insights, no study has systematically compared divergence in protein-coding sequences, gene expression profiles, and protein structures to assess variation across taxa or molecular levels. Two major challenges have historically hindered such analyses. First, experimental resolution of protein structures is time-consuming and technically demanding, leading to a scarcity of such data for many genes and most species [Slabinski et al., 2007, Jumper et al., 2021]. However, recent advances in artificial intelligence have enabled fast and accurate predictions of protein structures [Baek et al., 2021, AlQuraishi, 2021, Jumper et al., 2021, Pearce and Zhang, 2021, Varadi et al., 2022], effectively overcoming this limitation. Second, it is challenging to compare divergence at different molecular levels because they are quantified using distinct metrics that can vary by orders of magnitude [Duret and Mouchiroud, 2000, Tirosh and Barkai, 2008, Kalinka et al., 2010, Warnefors and Kaessmann, 2013]. To address this problem, we devised a novel standardized approach to facilitate direct comparisons of divergence across taxa and molecular levels. Using these standardized metrics, we investigated variation in evolutionary divergence across four taxa and three molecular levels, as well as functional differences between genes under strong or weak constraint on different molecular levels.

## Results

### Derivation of comparable evolutionary divergence metrics

Based on previous findings [Lemos et al., 2005, Jordan et al., 2005, Gillis and Pavlidis, 2009, Buschiazzo et al., 2012, Warnefors and Kaessmann, 2013, Konate et al., 2019, James et al., 2021, Lau et al., 2024], we hypothesized that protein-coding sequences, gene expression profiles, and protein structures are modified to different degrees during the course of genic evolution. Therefore, a central goal of our study was to assess differences in evolutionary divergence across these molecular levels. To address this problem, we first extracted pairs of 1:1 orthologs from four taxa: *Drosophila* (*D. melanogaster* and *D. pseudoobscura*), rodents (mouse and rat), primates (human and macaque), and *Arabidopsis* (*A. thaliana* and *A. lyrata*; see *Methods*). To estimate raw protein-coding sequence divergence, we computed the nonsynonymous substitution rate (*𝐾_𝑎_*) between protein-coding sequences of each pair of orthologs (see *Methods*), as this provides a measure of the protein-modifying changes in a gene sequence. To estimate raw gene expression divergence, we calculated the Euclidean distance between multi-tissue expression profiles of each pair of orthologs (see *Methods*), which is an ideal metric for expression divergence [Pereira et al., 2009, Glazko and Mushegian, 2010] that has been used in prior studies [Jordan et al., 2005, Yanai et al., 2004, Liao and Zhang, 2006, Warnefors and Kaessmann, 2013, Assis, 2019a]. To estimate raw protein structural divergence, we computed a similarity score between aligned protein structures of each pair of orthologs, which we converted to a difference representing protein structural divergence (see *Methods*).

Summary statistics of raw divergence metrics revealed substantial variation across molecular levels within each taxon (Tables S1–S4). To enable meaningful comparisons of divergence in protein-coding sequences, gene expression profiles, and protein structures, we standardized these metrics onto a common scale using a two-step approach (Figure 1). First, we applied Box-Cox transformations [Box and Cox, 1964] to normalize the raw distributions. We selected Box-Cox for its flexibility in estimating the optimal transformation parameter (*𝜆*) for each dataset, allowing it to accommodate different distribution shapes and ensure appropriate normalization. Second, we computed two sets of *𝑧*-scores to place the normalized divergence values on the same unitless scale. In particular, a *𝑧*-score quantifies how many standard deviations a value lies from the mean of its distribution, making it a useful metric for comparing otherwise incommensurate data. The first set, denoted as ‘taxon-specific *𝑧*-scores’, was generated by grouping genes within each taxon and standardizing divergence across molecular levels (Tables S5-S8). Each taxon-specific *𝑧*-score therefore represents how much the divergence of a gene deviates from the mean divergence across levels for a particular taxon. The second set, denoted as ‘level-specific *𝑧*-scores’, grouped genes by molecular level and standardized divergence across taxa (Tables S9-S12; see *Methods*). Each level-specific *𝑧*-score hence represents how much the divergence of a gene deviates from the mean divergence across taxa for a particular molecular level. Together, these two sets of *𝑧*-scores provided standardized, unitless measures of divergence in protein-coding sequences, gene expression profiles, and protein structures, facilitating direct comparisons across metrics that originally differed in scale and units.

**Figure 1:**
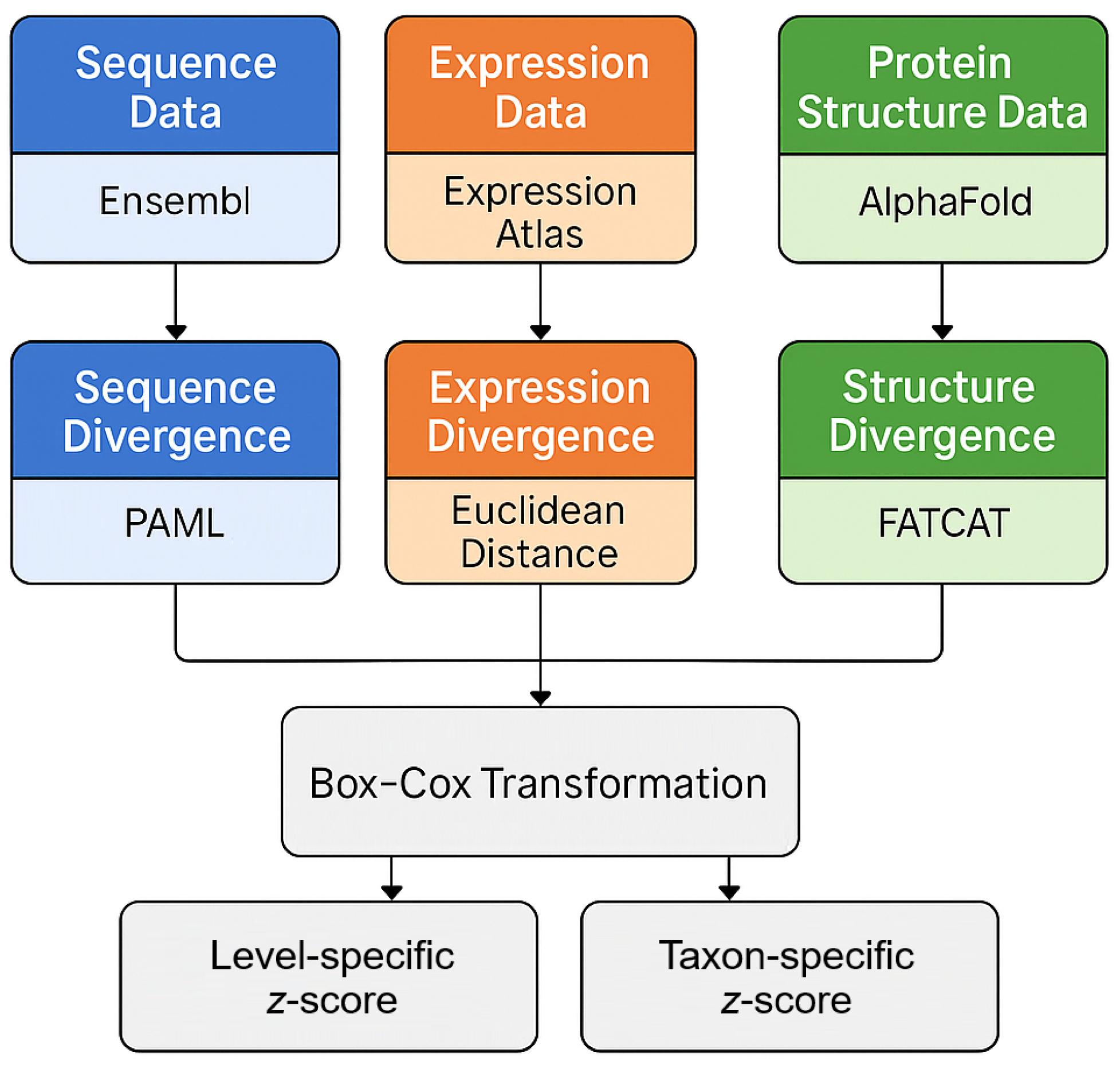
Schematic depicting the approach used to standardize raw divergence metrics.

### Comparisons of divergence across taxa and molecular levels

Our analysis of taxon-specific *𝑧*-scores uncovered striking differences in the divergence of protein-coding sequences, gene expression profiles, and protein structures among the four taxa (Figure 2). Specifically, sequence divergence was lowest in primates, followed by rodents, *Arabidopsis*, and *Drosophila* (Figure 2, left). This pattern aligns with theoretical expectations [Kimura and Ohta, 1971, Lynch and Conery, 2003] and with empirical findings from many sequence-based studies [Wu and Li, 1985, Sharp and Li, 1989, Martin and Palumbi, 1993, Lynch and Conery, 2003, Bromham and Penny, 2003, Figuet et al., 2015, Bromham, 2011, Lanfear et al., 2014], demonstrating the utility of our approach. In contrast, expression divergence was lowest in rodents, followed by *Arabidopsis*, primates, and *Drosophila* (Figure 2, middle), and this ranking was robust to the use of alternative expression datasets for rodents and primates (Figure S1, middle). Protein structural divergence was lowest in *Arabidopsis*, followed by rodents, primates, and *Drosophila* (Figure 2, right). These differences in ordering may be partially explained by the generally low-to-moderate associations between the evolution of protein-coding sequences and expression levels [Makova and Li, 2003, Nuzhdin et al., 2004, Lemos et al., 2005, Khaitovich et al., 2005, Assis and Kondrashov, 2014, Zhong et al., 2021], as well as between gene expression divergence and protein structural evolution [Bhardwaj and Lu, 2005, Paliy et al., 2008, Singh and Dash, 2008, Maier et al., 2009, Warnefors and Kaessmann, 2013]. Moreover, while protein structure is dependent on the coding sequence [Anfinsen, 1973], gene expression is often regulated by noncoding regions [Carroll, 2008, Wittkopp and Kalay, 2012, Consortium et al., 2012] and may therefore evolve at a different rate. Together, these results illustrate that the relative ordering of divergence across taxa depends strongly on the molecular level considered.

**Figure 2:**
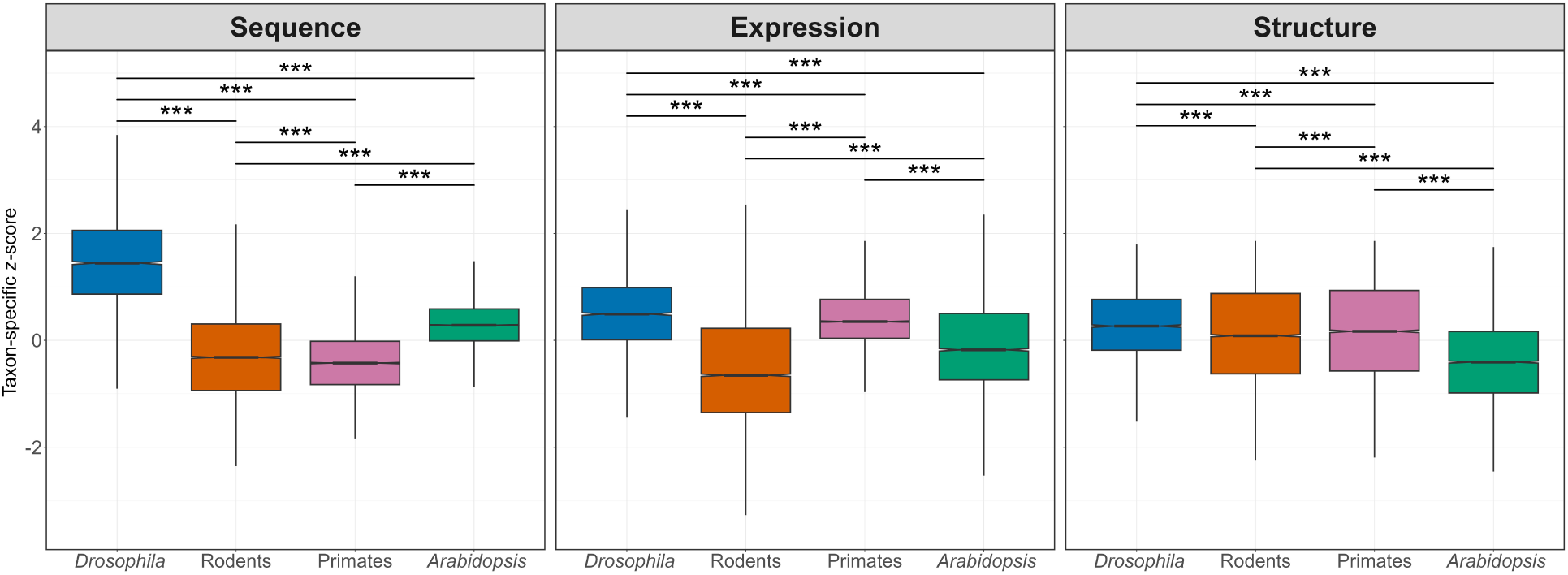
Comparisons of evolutionary divergence across taxa. Boxplots depict distributions of taxon-specific *𝑧*-scores for protein-coding sequences, gene expression profiles, and protein structures in *Drosophila*, rodents, primates, and *Arabidopsis*. \**𝑃 <* 0.05, \*\**𝑃 <* 0.01, \*\*\**𝑃* < 0.001 (see *Methods*).

Comparisons of level-specific *𝑧*-scores revealed a clear pattern in the divergence of protein-coding sequences, gene expression profiles, and protein structures within taxa (Figure 3). Across all four taxa, sequence divergence was lowest and protein structural divergence highest, and this pattern was consistent when we repeated our analysis using alternative gene expression data (Figure S2). This ordering is generally consistent with the idea that higher levels of biological organization are closer proxies for the functions on which selection acts. In particular, expression profiles are often considered better proxies for function than gene sequences because they represent the relative abundances of transcripts across different spatial (e.g., tissues) or temporal (e.g., developmental stages) conditions [Carroll, 2005, Nehrt et al., 2011, Assis and Bachtrog, 2013, De Smet et al., 2017], and protein structures may represent even better proxies due to their direct roles in cellular and intracellular processes [Chothia and Lesk, 1986, Petsko and Ringe, 2004]. Hence, these findings reveal a conserved hierarchy of divergence across molecular levels within taxa.

**Figure 3:**
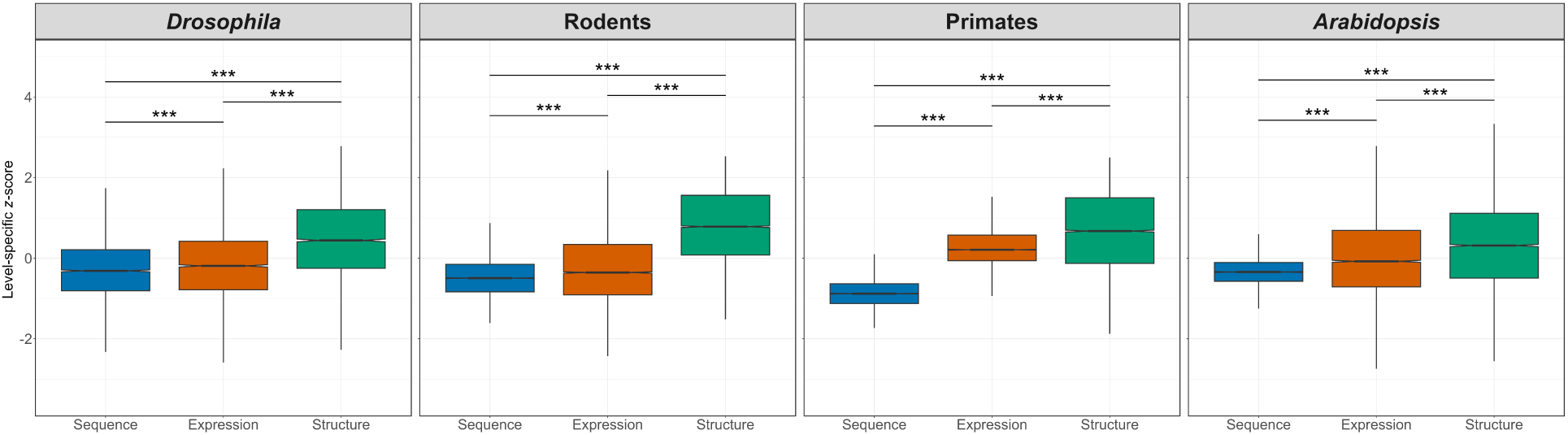
Comparisons of evolutionary divergence across molecular levels. Boxplots depict distributions level-specific *𝑧*-scores for protein-coding sequences, gene expression profiles, and protein structures in *Drosophila*, rodents, primates, and *Arabidopsis*. (see *Methods*).

### Assessment of relationships in divergence among molecular levels

Next, we sought to understand connections between evolutionary changes at the protein-coding sequence, gene expression, and protein structural levels. To address this question, we used taxon-specific *𝑧*-scores to evaluate relationships between divergence of coding sequences and expression profiles, coding sequences and protein structures, and expression profiles and protein structures in primates, rodents, *Drosophila*, and *Arabidopsis* (Figure 4). Across all four taxa, we observed small positive correlations of similar magnitudes between coding sequence and expression divergence (Figure 4, top row). These correlations remained consistent when raw divergence metrics were used instead of standardized scores (Figure S3). Similar patterns have been observed in many other studies [Makova and Li, 2003, Nuzhdin et al., 2004, Khaitovich et al., 2005, Lemos et al.,

**Figure 4:**
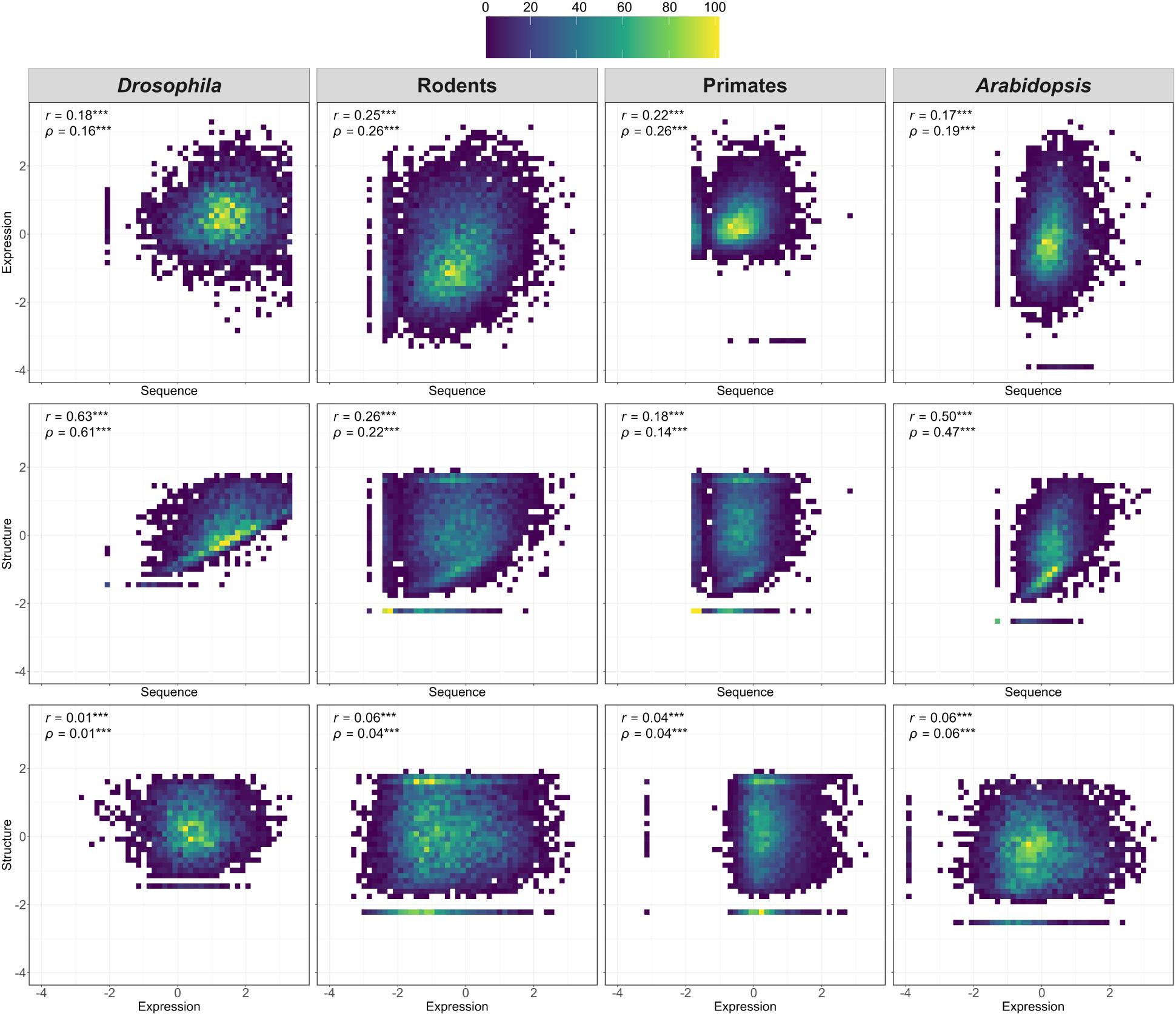
Relationships between evolutionary divergence of protein-coding sequences, gene expression profiles, and protein structures. Heatmaps depicting correlations between taxons-specific *𝑧*-scores of sequence and expression divergence (top row), sequence and structural divergence (middle row), and expression and structural divergence (bottom row) in *Drosophila*, rodents, primates, and *Arabidopsis*. Pearson (*𝑟*) and Spearman (*𝜌*) correlation coefficients are shown in the upper left corner of each plot. \**𝑃 <* 0.05, \*\**𝑃* < 0.01, ***𝑃 < 0.001 (see *Methods*).

2005, Kalinka et al., 2010, Assis and Kondrashov, 2014, Harnqvist et al., 2021, Zhong et al., 2021], supporting the widely held view that alterations in both protein-coding and regulatory sequences contribute to gene expression divergence, with regulatory sequences outside of the coding region expected to play a greater role [Wray, 2007, Carroll, 2008, Wittkopp and Kalay, 2012]. Additionally, this imperfect relationship may help explain the significant differences between rates of sequence and expression divergence observed in all four taxa (Figure 3).

Similarly, coding sequence and structural divergence were positively correlated in all four taxa (Figure 4, middle row), consistent with prior studies [Sanchez and Sali, 1998, Wood and Pearson, 1999, Tirosh and Barkai, 2008, Marks et al., 2012, Kc, 2017] and the known dependency of protein structures on underlying coding sequences [Anfinsen, 1973]. However, while the relationship between sequence and structural divergence was moderately strong in *Drosophila* and *Arabidopsis*, it was quite weak in rodents and primates, indicating that nonsynonymous substitutions contribute less to the evolution of protein structures in these taxa. In particular, nonsynonymous substitutions can have varying effects on protein structure depending on their chemical properties, such as composition, polarity, and molecular volume [Grantham, 1974, Yampolsky and Stoltzfus, 2005]. Regardless of the underlying reason, this finding points to some uncoupling between sequence and structural evolution, suggesting that genes may have different targets of selection.

Our comparisons of gene expression and structural divergence uncovered very weak positive relationships in all taxa (Figure 4, bottom row)—much weaker than the other observed relationships. This finding is compatible with the idea that expression is typically regulated by regions outside of the coding region of a gene [Carroll, 2008, Wittkopp and Kalay, 2012, Consortium et al., 2012]. Although statistically significant, the correlations were only slightly greater than zero in primates and rodents, possibly reflecting taxon-specific differences in the relationship between expression and structural evolution. Prior studies have similarly found inconsistent relationships between expression and structural divergence, with some reporting weak positive correlations and others weak negative correlations [Bhardwaj and Lu, 2005, Paliy et al., 2008, Singh and Dash, 2008, Maier et al., 2009, Warnefors and Kaessmann, 2013].

An alternative possibility for the lack of observed correlations with protein structural divergence is that relationships may be multivariate. To explore potentially more complex interactions among divergence of protein-coding sequences, gene expression profiles, and protein structures, we fit a set of linear models to predict one of the three divergence metrics (the response variable) from the other two (feature variables; see *Methods*). Specifically, we fit models to predict coding sequence divergence (response) from gene expression and protein structural divergence (features), gene expression divergence (response) from coding sequence and protein structural divergence (features), and protein structural divergence (response) from coding sequence and gene expression divergence (features). Regardless of the chosen response variable, the best-fit models failed to uncover new relationships between divergence at different molecular levels in any of the four taxa (Table S13). Across all taxa, the only statistically significant relationships identified were between sequence and expression divergence and between sequence and protein structural divergence. Thus, there do not appear to be direct relationships between gene expression and protein structural evolution. Taken together, these comparisons support the hypothesis that evolutionary targets vary across genes, with distinct sets of genes undergoing rapid sequence, expression, and structural divergence.

### Characterization of divergence at different molecular levels

Last, we considered the possibility that evolutionary targets vary across genes based on their functions. For example, if a gene encodes a protein with critical binding properties, then its structure may be under stronger constraint than its sequence or expression profile. To test this hypothesis, we conducted gene ontology (GO) enrichment analyses of genes exhibiting low (bottom 2.5%) and high (top 2.5%) divergence in protein-coding sequences, gene expression profiles, and protein structures (see *Methods*). We found that genes with low sequence divergence were enriched for functions related to protein synthesis and transport in all taxa (Tables S14-17), and specifically for neuronal developmental functions in mammals. In contrast, genes with high sequence divergence were often involved in immune-related functions in all taxa (Tables S18–S21). These patterns align with previous findings linking slow sequence divergence to essential biological processes [Green, 1941, Dudek et al., 1997, Chan et al., 2009, Liao et al., 2014, Li et al., 2016] and rapid sequence divergence to immune response [van Loon et al., 2006, Barreiro and Quintana-Murci, 2010, Liao et al., 2014, Grueber et al., 2014, Assis, 2019a]. In particular, the rapid sequence divergence of immune-related genes may be driven, at least in part, by the need to adapt to emerging pathogens [Litman et al., 2005, Dzik, 2010].

Examining expression divergence, we found a similar dichotomy. Genes with low expression divergence were primarily involved in maintaining cellular homeostasis, with functions such as energy production, protein synthesis, and protein transport (Tables S22–S25), underscoring the necessity of conserving essential biological processes. Conversely, genes with high expression divergence were enriched for roles in signal transduction and sexual reproduction, with some taxon-specific specializations, such as neuronal signaling in mammals and response to stimuli in *Arabidopsis* (Tables S26–S29) [Zou et al., 2009]. Notably, the most enriched GO term in primates was ‘xenobiotic metabolic process’, which governs the breakdown of foreign chemicals, such as drugs, pesticides, and environmental contaminants. This suggests that while core cellular functions remain tightly constrained, regulatory mechanisms governing external interactions are more evolutionarily flexible to promote adaptability in dynamic environments [Espinosa-Soto et al., 2011, Babbitt et al., 2017, Jovanovic et al., 2021, Couce, 2024, Durkin et al., 2024].

Extending this analysis to protein structure, we found that genes with low structural divergence were predominantly associated with protein synthesis, processing, binding, and transport (Tables S30–S33), reflecting strong purifying selection on proteins essential for cellular integrity. In contrast, genes with high structural divergence frequently encode proteins involved in membrane transport, gene regulation, and organelle-specific functions, particularly within the nucleus (Tables S34-S37). These findings suggest that while the structural stability of translation and transport proteins is critical for cellular function, regulatory and membrane-associated proteins may undergo greater structural diversification to accommodate lineage-specific adaptations. This pattern mirrors broader evolutionary trends, where highly conserved proteins are functionally indispensable, while structurally dynamic proteins facilitate adaptive responses to environmental challenges [Drummond et al., 2005, Rorick and Wagner, 2010, Smith and Workman, 2012]. For example, GTP-related functions were highly enriched among genes with low structural divergence in all taxa, reflecting their conserved roles in cellular regulation and molecular transport [Bourne et al., 1990, Pfeffer, 1992, Ma, 2007]. Conversely, genes encoding mRNA-binding proteins often exhibited high structural divergence, perhaps due to the flexibility required for post-transcriptional regulation [Lunde et al., 2007, Ma et al., 2023].

To delve more deeply into differences in evolutionary targets across genes, we examined overlaps among sets of genes with low and high divergence in their sequences, expression profiles, and protein structures in each taxon (Figure 5; Table S38). This analysis revealed minimal overlap among gene sets, reinforcing the idea that selection can act independently on gene sequences, expression profiles, and protein structures. As a baseline for comparison, we generated randomly sampled gene sets and found no overlap in any taxon, demonstrating that the limited overlap observed in the empirical data still exceeds random expectation. Specifically, among sets with low divergence, there were zero common genes in *Drosophila*, three in rodents, zero in primates, and two in *Arabidopsis*. As expected, these conserved genes are involved in core cellular functions, such as transcriptional and translational regulation, signal transduction, and transmembrane transport. Similarly, among sets with high divergence, there was one common gene in *Drosophila*, zero in rodents, two in primates, and five in *Arabidopsis*. Notably, *SMIM24* (ENSG00000095932) in primates and *MTO3* (AT3G17390) in *Arabidopsis* both function in pathogen defense through their roles in cell membrane processes. Taken together with our broader analyses, these findings highlight how evolutionary pressures may act through distinct mechanisms on sequence, expression, and structural divergence, shaping genetic diversity across taxa.

**Figure 5:**
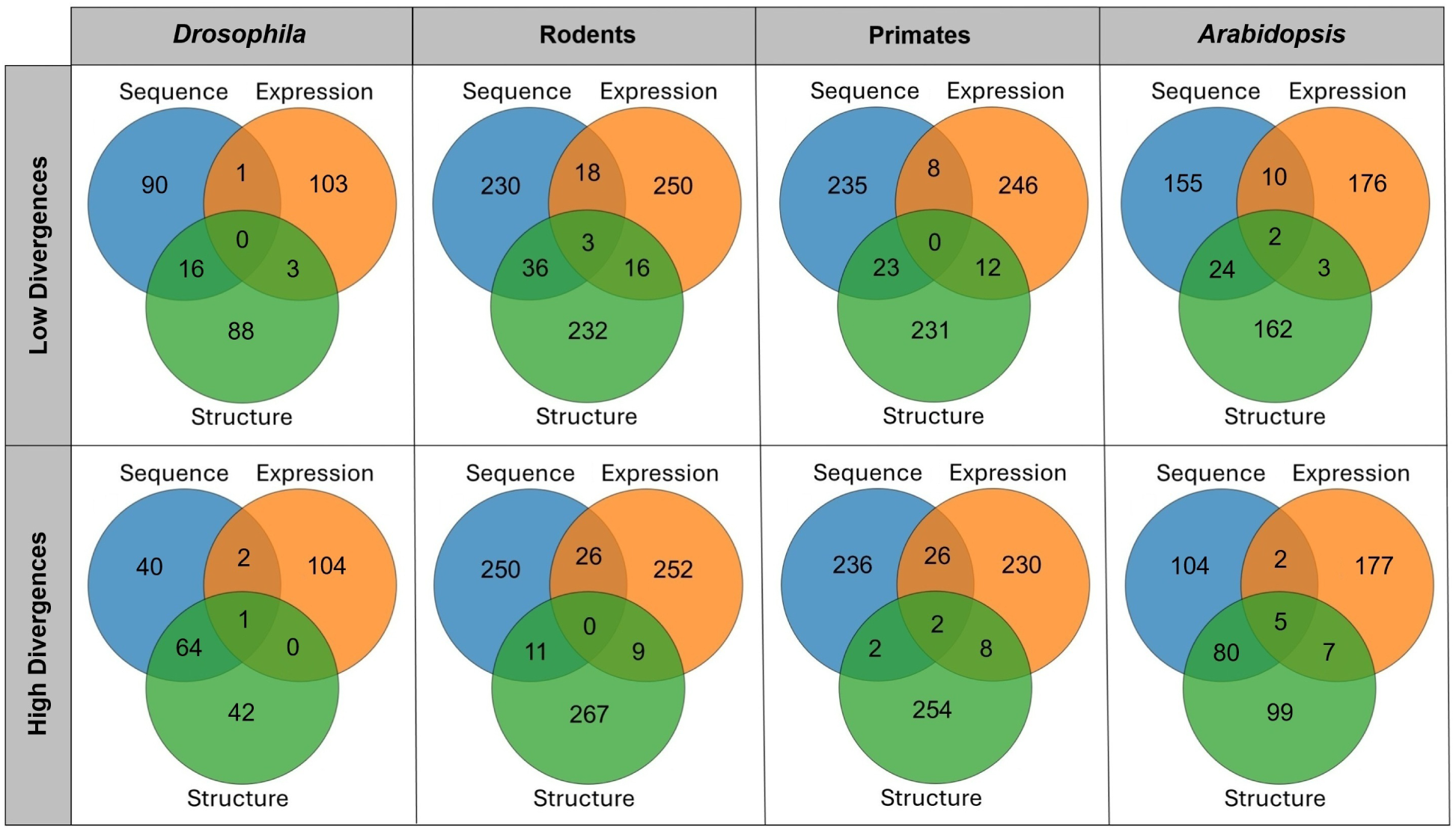
Overlaps among gene sets with low or high divergence of their protein-coding sequences, expression profiles, and protein structures. Venn diagrams showing numbers of common genes with low (top row) and high (bottom row) sequence, expression, and structural divergence in *Drosophila*, rodents, primates, and *Arabidopsis*; see *Methods*).

## Discussion

This study introduces a standardized approach for directly comparing divergence in protein-coding sequences, gene expression profiles, and protein structures across taxa using *𝑧*-scores. While previous research has examined divergence across different taxa and molecular levels [Sanchez and Sali, 1998, Duret and Mouchiroud, 2000, Makova and Li, 2003, Choi and Kim, 2006, Paliy et al., 2008, Buschiazzo et al., 2012, Warnefors and Kaessmann, 2013, Gillis and Pavlidis, 2009], our method is the first to integrate three classes of divergence within a unified framework, facilitating a more comprehensive assessment of evolutionary dynamics. By standardizing divergence, we ensured comparability among taxa and molecular levels, overcoming limitations of previous analyses. While we applied this approach to *Drosophila*, rodents, primates, and *Arabidopsis*, its design allows for broad applicability across any taxa, making it a valuable tool for future research in evolutionary genomics.

Our comparisons of evolutionary divergence across taxa revealed that protein-coding sequences, gene expression profiles, and protein structures all evolve fastest in *Drosophila*, aligning with theoretical predictions based on effective population size (*𝑁_𝑒_*) and mutation rate (*𝜇*) [Kimura and Ohta, 1971, Lynch and Conery, 2003] and with previous empirical findings of rapid evolution in this taxon [Britten, 1986, Moriyama, 1987, Sharp and Li, 1989, Lynch and Conery, 2003, Bromham, 2011, Coolon et al., 2014, Nourmohammad et al., 2017]. Indeed, using the synonymous substitution rate (*𝐾_𝑠_*) as a crude estimate of mutation rate also supports this pattern (Figure S4). However, while *Drosophila* consistently exhibited the fastest evolution across all molecular levels, the relative rankings of the other three taxa varied depending on the level examined. Despite these differences across taxa, our comparisons of evolutionary divergence across molecular levels demonstrated that structural divergence was fastest in all taxa, supporting the notion that protein structures most closely reflect the functions on which selection acts. Moreover, nearly all taxon-specific relationships among different classes of divergence were weak, pointing to limited interactions between evolutionary processes at different molecular levels. Together, these findings highlight the taxon-specific variability of selection acting on different levels of biological organization, suggesting that evolutionary targets vary across genes.

Building on these findings, we hypothesized that evolutionary targets of genes depend on their functions. Supporting this hypothesis, we identified distinct functional enrichments for genes with slow and fast divergence at different molecular levels. While genes with slow divergence generally contribute to core biological functions, we also observed some clear differences across molecular levels. Specifically, slow sequence divergence was associated with developmental processes related to transcription and neuronal differentiation, slow expression divergence with protein homeostasis and mitochondrial activity, and slow structural divergence with essential cellular machinery for protein synthesis, including ribosomes, spliceosomes, and proteasomes. Similarly, functions of genes with fast divergence at different levels also differed greatly. Fast sequence divergence was associated with immune-related processes such as antiviral defense and T-cell signaling, fast expression divergence with extracellular signaling and ion transport, and fast structural divergence with intracellular regulation, including chromatin remodeling, organelle dynamics, and transcriptional control. Further analysis showed minimal overlap among gene sets with low or high divergence at different molecular levels, implying that selection may allow changes at one level while maintaining stability in others. Collectively, these findings suggest that evolutionary pressures may shape sequence, expression, and structural divergence in function-specific ways across taxa.

## Methods

### Data acquisition and processing

Sequence data for this study were obtained from Ensembl databases [Dyer et al., 2025]. *Drosophilia* sequences (*D. melanogaster* and *D. pseudoobscura*) were retrieved from https://metazoa.ensembl.org/, mammalian sequences (*Mus musculus*, *Rattus norvegicus*, *Homo sapiens*, and *Macaca mulatta*) from https://www.ensembl.org/, and *Arabidopsis* sequences (*A. thaliana* and *A. lyrata*) from https://plants.ensembl.org/. For each of the four species pairs (*D. melanogaster* –*D. pseudoobscura*, mouse–rat, human–macaque, and *A. thaliana*–*A. lyrata*), 1:1 orthologous protein-coding genes were obtained from Ensembl BioMart at[Dyer et al., 2025], which infers orthology using sequence similarity, gene family clustering, and phylogenetic tree reconstruction to distinguish speciation from duplication events. For mammals, we downloaded multiple alignments of exons for all protein-coding genes from the UCSC Genome Browser [Kent et al., 2002, Nassar et al., 2023] at https://genome.ucsc.edu/, concatenated aligned exons by gene, and extracted those corresponding to the four species considered here. Coding sequences for *Drosophila* and *Arabidopsis* were downloaded and aligned on a gene-by-gene basis with MACSE [Ranwez et al., 2018]. The resulting pairwise alignments were then analyzed using the codeml program in the PAML package [Yang, 2007] to estimate the nonsynonymous substitution rate (*𝐾_𝑎_*) and synonymous substitution rate (*𝐾_𝑠_*) for each orthologous gene pair.

Expression data consisted of tables of quantile-normalized RNA-seq abundances computed in transcripts per million (TPM) from six tissues in *Drosophila* [Assis, 2019b], nine tissues in rodents (E-MTAB-3718 and E-MTAB-2800), six tissues in primates (E-MTAB-3716 and E-MTAB-3717), and three tissues in *Arabidopsis* (E-GEOD-38612 and E-MTAB-5072). We downloaded these data from the Dryad digital repository at https://www.datadryad.org [Vision, 2010] for *Drosophila* and the Gene Expression Atlas [Kapushesky et al., 2010] at https://www.ebi.ac.uk/gxa/home/ for mammals and *Arabidopsis*. The six tissues in the *Drosophila* dataset are carcass, female head, male head, ovary, accessory gland, and testis [Assis, 2019b]. The nine tissues in the rodent dataset are brain, heart, kidney, liver, testis, lung, spleen, colon, and skeletal muscle [Merkin et al., 2012]. The six tissues in the primate dataset are brain, cerebellum, heart, kidney, liver, and testis [Brawand et al., 2011]. The three tissues in the *Arabidopsis* dataset are flower, seed, and root [Haudry et al., 2013]. To assess the robustness of our method, we analyzed two additional independently generated gene expression datasets, one from rodents (E-GEOD-74747 and E-GEOD-53960) [Yu et al., 2014, Huntley et al., 2016] and one from primates (E-MTAB-4344 and E-MTAB-2799) [Merkin et al., 2012, Lin et al., 2014]. To minimize noise, we removed lowly expressed genes, or those for which TPM *<* 1 in all tissues. All remaining TPM values were log-transformed and relativized across tissues to enable comparisons between species.

Protein structure data were downloaded from the AlphaFold database [Jumper et al., 2021] at https://alphafold.ebi.ac.uk/ in Protein Data Bank (PDB) file format. Each PDB file contains the three-dimensional atomic coordinates of a protein. PDB files of each pair of 1:1 orthologs were provided as input to FATCAT version 2.0 [Li et al., 2020], which performs a series of superpositions of the protein structures onto each other and computes a *𝑝*-value, chaining score, root mean square deviation (RMSD), number of twists needed for better superposition (*𝑛*), and total number of equivalent superposed structures (*𝐿*). FATCAT uses these values to estimate a similarity score *𝑠* ranging from 0 to 100%.

### Divergence estimations and comparisons

We combined processed sequence, expression, and protein structure datasets for each taxon. After removing genes with missing data, we obtained final datasets consisting of 4,277 1:1 orthologs in *Drosophila*, 11,483 1:1 orthologs in rodents, 10,668 1:1 orthologs in primates, and 7628 1:1 orthologs in *Arabidopsis*. For each pair of 1:1 orthologs, we used the computed *𝐾_𝑎_* value as an estimate of raw protein-coding sequence divergence, the Euclidean distance between relativized expression values as an estimate of raw gene expression divergence, and the percentage difference between aligned structures (100 − *𝑠*) as an estimate of raw protein structural divergence.

To enable comparisons among these divergence metrics, we first normalized their distributions by applying Box-Cox transformations [Box and Cox, 1964] with the SciPy library in Python [Virtanen et al., 2020]. Then, we standardized the normalized divergence metrics by converting them to two sets of *𝑧*-scores. In particular, for comparisons among taxa (i.e., primates vs. rodents vs. grasses), we computed taxon-specific *𝑧*-scores to measure distances between each divergence of a gene and the mean across levels for a particular taxon. In contrast, for comparisons among divergence at different molecular levels (i.e., sequence vs. expression vs. structural), we computed level-specific *𝑧*-scores to measure distances between each divergence of a gene and the mean across taxa for a particular level.

### Statistical analyses

Functional enrichment analysis was performed with the DAVID Functional Annotation Tool [Sherman et al., 2022] at https://david.ncifcrf.gov/tools.jsp (last accessed on December 15, 2025). In particular, we generated a ranked list of genes for each species and molecular level, extracted the top or bottom 2.5% of genes, and evaluated functional enrichment of these genes against the genome-wide background. We then obtained Gene Ontology (GO) [Ashburner et al., 2000] terms describing enriched cellular components, biological processes, or molecular functions of genes in each list. To account for multiple testing, we only considered a GO term as significantly enriched if *𝑃 <* 0.05 after Benjamini-Hochberg adjustment for false discovery rate [Benjamini and Hochberg, 1995].

All other statistical analyses were performed in the R software environment [R Core Team, 2021]. Two-tailed Mann-Whitney *𝑈* tests implemented with the geom signif() function in the package ggsignif [Ahlmann-Eltze and Patil, 2021] were used to compare all pairs of distributions shown in Figures 2, 3, S1, S2, and S5. We used two approaches to evaluate relationships between or among divergence at different molecular levels. First, we evaluated pairwise relationships by computing Pearson (*𝑟*) and Spearman (*𝜌*) correlation coefficients between different classes of divergence with the cor.test() function in the stats package [R Core Team, 2021]. Second, we assessed more complex relationships among the three classes of divergence by fitting multiple linear regression models with the lm() function in the stats package [R Core Team, 2021]. For each taxon, we fit three models, in which each model used a different divergence as the predicted response and the other two divergence metrics as the input features.

## Data availability

All data supporting the findings of this study are available within the paper and its Supplementary Information.

## Acknowledgments

This work was supported by National Institutes of Health grant R35GM142438 and National Science Foundation grant DBI-2130666.

