## Supplementary material for "Gene function predicts divergence across molecular levels": Figures S1-S4

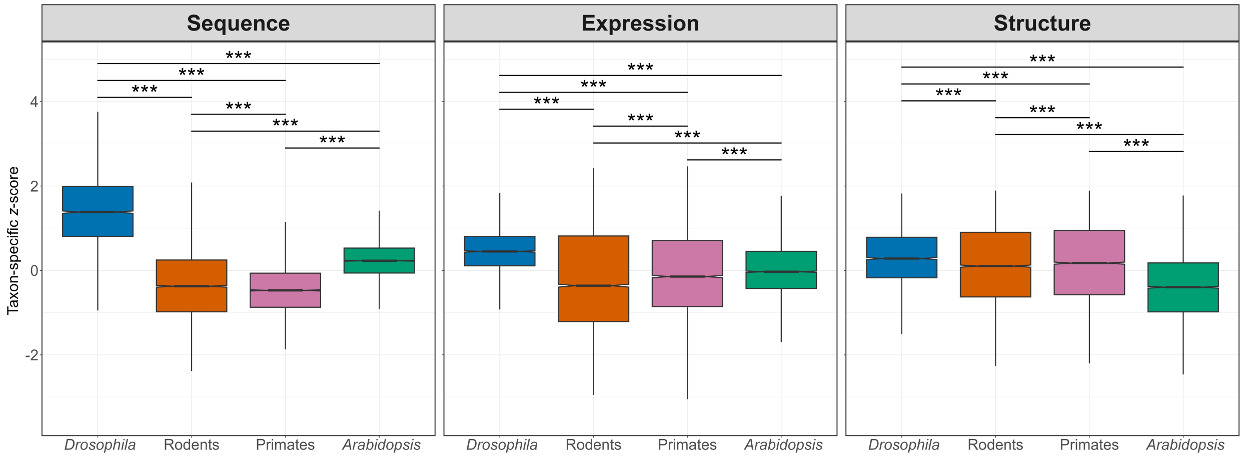


Figure S1: Comparisons of evolutionary divergence across taxa. Boxplots depict distributions of taxon-specific $z$-scores for protein-coding sequences, gene expression profiles (different dataset than Figure 2 for rodents and primates), and protein structures in *Drosophila*, rodents, primates, and *Arabidopsis*. $^*P < 0.05$, $^{**}P < 0.01$, $^{***}P < 0.001$ (see *Methods*).

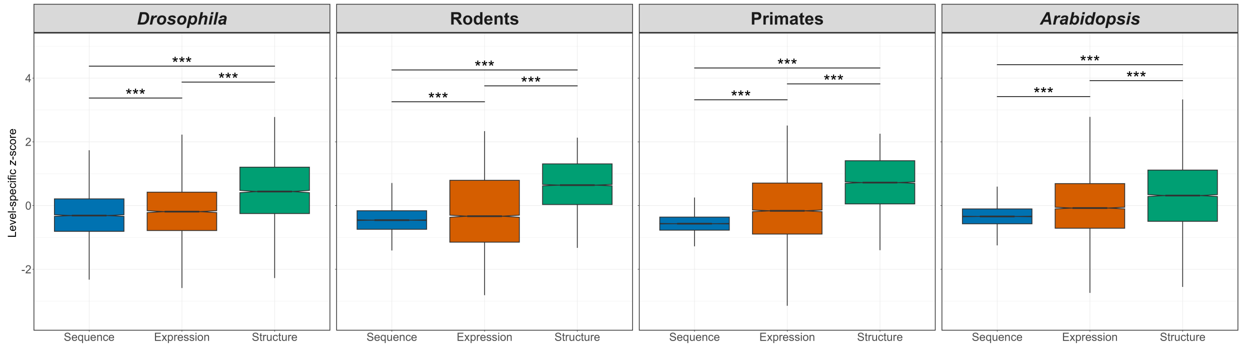


Figure S2: Comparisons of evolutionary divergence across molecular levels. Boxplots depict distributions of level-specific $z$-scores for protein-coding sequences, gene expression profiles (different dataset than Figure 2 for rodents and primates), and protein structures in *Drosophila*, rodents, primates, and *Arabidopsis*. $^*P < 0.05$, $^{**}P < 0.01$, $^{***}P < 0.001$ (see *Methods*).

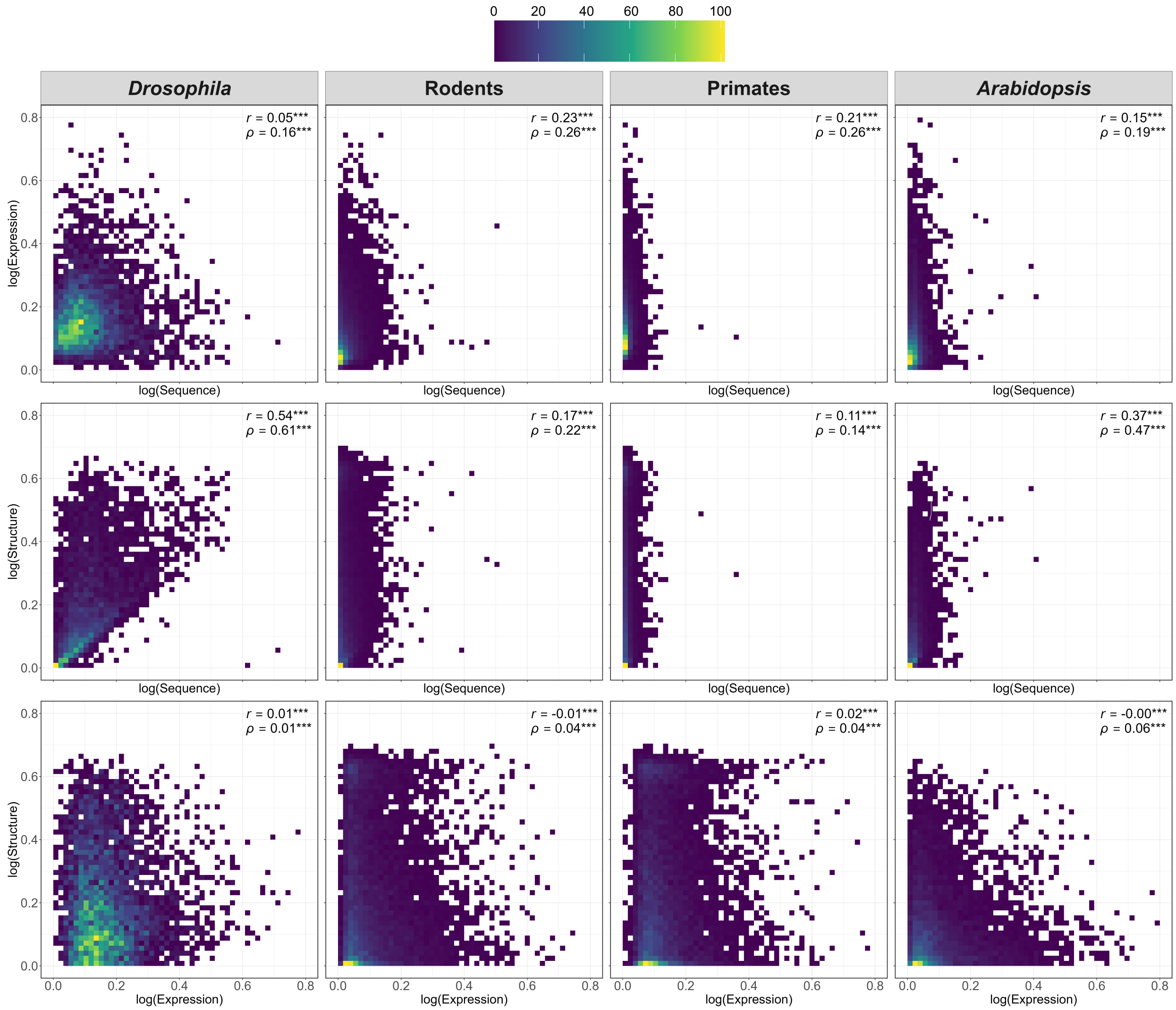


Figure S3: Relationships between raw evolutionary divergence of protein-coding sequences, gene expression profiles, and protein structures. Heatmaps depicting correlations between raw sequence and expression divergence (top row), sequence and structural divergence (middle row), and expression and structural divergence (bottom row) in *Drosophila*, rodents, primates, and *Arabidopsis*. Pearson ($r$) and Spearman ($\rho$) correlation coefficients are shown in the upper right corner of each plot. $^{*}P < 0.05$, $^{**}P < 0.01$, $^{***}P < 0.001$ (see *Methods*).

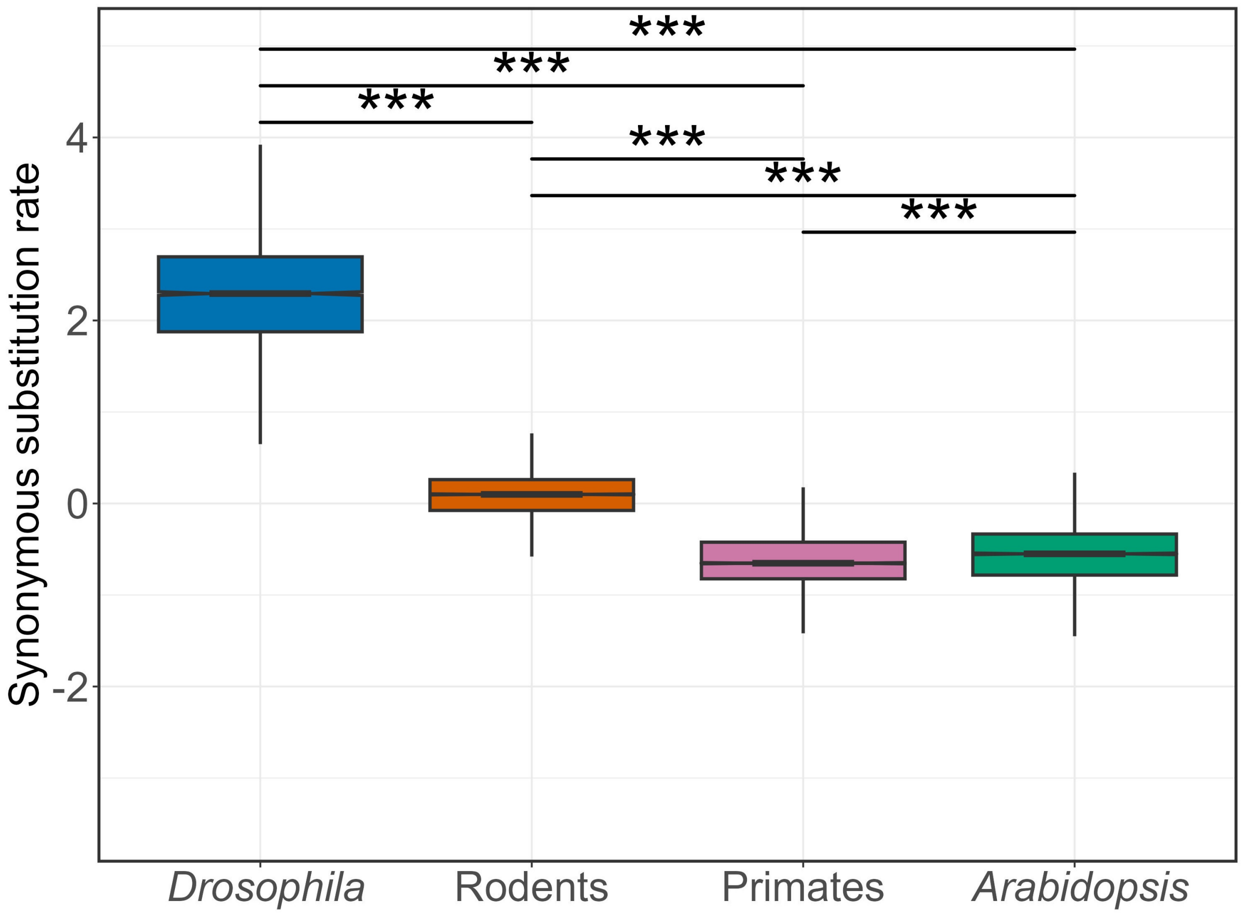


Figure S4: Comparison of synonymous substitution rates ($K_s$) among taxa. Boxplots depict distributions of $K_s$ in primates, rodents, drosophilids, and arabidopsis. *$P < 0.05$, **$P < 0.01$, ***$P < 0.001$ (see *Methods*).
